# Joint coding of visual input and eye/head position in V1 of freely moving mice

**DOI:** 10.1101/2022.02.01.478733

**Authors:** Philip R. L. Parker, Elliott T. T. Abe, Emmalyn S. P. Leonard, Dylan M. Martins, Cristopher M. Niell

**Author notes:** Authors contributed equally (Philip R. L. Parker, Elliott T. T. Abe).

## Abstract

**SUMMARY:** Visual input to the brain during natural behavior is highly dependent on movements of the eyes, head, and body. Neurons in mouse primary visual cortex (V1) respond to eye and head movements, but how information about eye and head position is integrated with visual processing during free movement is unknown, since visual physiology is generally performed under head-fixation. To address this, we performed single-unit electrophysiology in V1 of freely moving mice while simultaneously measuring the mouse’s eye position, head orientation, and the visual scene from the mouse’s perspective. Based on these measures we were able to map spatiotemporal receptive fields during free movement, using a generalized linear model (GLM) that predicted the activity of V1 neurons based on gaze-corrected visual input. Furthermore, we found that a significant fraction of visually-responsive neurons showed tuning for eye position and head orientation. Incorporating these variables into the GLM revealed that visual and positional signals are integrated through a multiplicative mechanism in the majority of modulated neurons, consistent with computation via gain fields and nonlinear mixed selectivity. These results provide new insight into coding in mouse V1, and more generally provide a paradigm for performing visual physiology under natural conditions, including active sensing and ethological behavior.

**HIGHLIGHTS:**

- Neurons in mouse V1 respond to both vision and self-motion, but it is unclear how these are combined.
- We record neural activity in V1 concurrent with measurement of the visual input from the mouse’s perspective during free movement.
- These data provide the first measurement of visual receptive fields in freely moving animals.
- We show that many V1 neurons are tuned to eye position and head orientation, and these contribute a multiplicative gain on visual responses in the majority of modulated neurons.

## INTRODUCTION

A key aspect of natural behavior is movement through the environment, which has profound impacts on the incoming sensory information (Gibson, 1979). In vision, movements of the eyes and head due to locomotion and orienting transform the visual scene in ways that are potentially both beneficial, by providing additional visual cues, and detrimental, by introducing confounds due to self-movement. By accounting for movement, the brain can therefore extract more complete and robust information to guide visual perception and behavior. Accordingly, a number of studies have demonstrated the impact of movement on activity in cortex (Busse et al., 2017; Froudarakis et al., 2019; Parker et al., 2020). In head-fixed mice, locomotion on a treadmill increases the gain of visual responses (Niell and Stryker, 2010) and modifies spatial integration (Ayaz et al., 2013) in V1, while passive rotation generates vestibular signals (Bouvier et al., 2020; Vélez-Fort et al., 2018). Likewise, in freely moving mice and rats, V1 neurons show robust responses to head and eye movements and head orientation tuning (Guitchounts et al., 2020a, 2020b; Meyer et al., 2018).

However, it is unknown how information about eye and head position is integrated into visual processing during natural movement, since studies of visual processing are generally performed during head-fixation to allow presentation of controlled stimuli, while natural eye and head movements require a mouse to be freely moving. Quantifying visual coding in freely moving animals requires determining the visual input, which is no longer under the experimenter’s control and is dependent on both the visual scene from the mouse’s perspective and its eye position. In addition, natural scenes, particularly during free movement, pose difficulties for data analysis since they contain strong spatial and temporal correlations and are not uniformly sampled because they are under the animal’s control.

To address the experimental challenge, we combined high density silicon probe recordings with miniature head-mounted cameras (Meyer et al., 2018; Michaiel et al., 2020; Sattler and Wehr, 2021), with one camera aimed outwards to capture the visual scene from the mouse’s perspective (“world camera”), and a second camera aimed at the eye to measure pupil position (“eye camera”), as well as an inertial measurement unit (IMU) to quantify head orientation. To address the data analysis challenge, we implemented a paradigm to correct the world camera video based on measured eye movements with a shifter network (Walker et al., 2019; Yates et al., 2021) and then use this as input to a generalized linear model (GLM) to predict neural activity (Pillow et al., 2008).

Using this approach, we first quantified the visual encoding alone during free movement, in terms of linear spatiotemporal receptive fields (RFs) from the GLM fit. For many units, the RF measured during free movement is similar to the RF measured with standard white noise stimuli during head-fixation within the same experiment, providing confirmation of this approach. We then extended the encoding model to incorporate eye position and head orientation, and found that these generally provide a multiplicative gain on the visual response. Together, this work provides new insights into the mechanisms of visual coding in V1 during natural movement, and opens the door to studying the neural basis of behavior under ethological conditions.

## RESULTS

### Visual physiology in freely moving mice

In order to study how visual processing in V1 incorporates self-motion, we developed a system to perform visual physiology in freely moving mice (Figure 1A). To estimate the visual input reaching the retina, a forward-facing world camera recorded a portion (~120 deg) of the visual scene available to the right eye. A second miniature camera aimed at the right eye measured pupil position, and an IMU tracked head orientation. Finally, a driveable linear silicon probe implanted in left V1 recorded the activity of up to 100+ single units across layers. The same neurons were first recorded under head-fixed conditions to perform white noise RF mapping, and then under conditions of free movement (Figure 1B). Well isolated units were highly stable across the two conditions (Figure S1 and Methods). Figure 1C and Video S1 show example data obtained using this system in a freely moving animal. Mice were allowed to explore a visually complex arena containing black and white blocks (three-dimensional sparse noise), static white noise and oriented gratings on the walls, and a monitor displaying moving spots. After several days of habituation, mice were active for a majority of the time spent in the arena (82%), with an average movement speed of 2.6 cm/s, which is comparable to other similar studies (see Methods; (Juavinett et al., 2019; Meyer et al., 2018).

**Figure 1:**
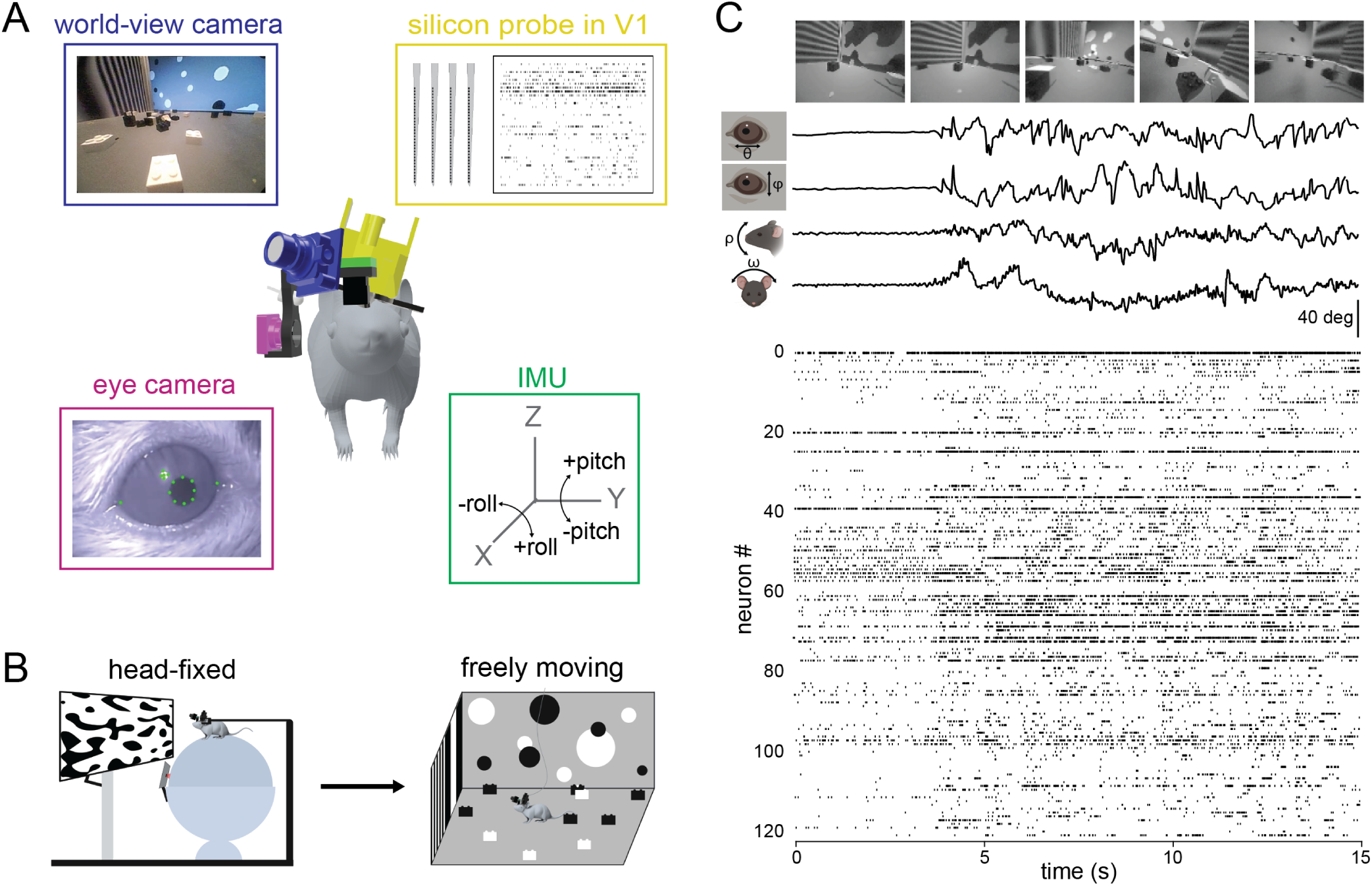
Visual physiology in freely moving mice. **A)** Schematic of recording preparation including 128-channel linear silicon probe for electrophysiological recording in V1 (yellow), miniature cameras for recording the mouse’s eye position (magenta) and visual scene (blue), and inertial measurement unit for measuring head orientation (green). **B)** Experimental design: controlled visual stimuli were first presented to the animal while head-fixed, then the same neurons were recorded under conditions of free movement. **C)** Sample data from a fifteen second period during free movement showing (from top) visual scene, horizontal and vertical eye position, head pitch and roll, and a raster plot of over 100 units. Note that the animal began moving at ~4 secs, accompanied by a shift in the dynamics of neural activity.

### A generalized linear model accurately estimates spatiotemporal receptive fields during free movement

To quantify visual coding during free movement, both the neural activity and the corresponding visual input are needed. The world camera captures the visual scene in a head-centric point of view, while the visual input needed is in a retinocentric perspective. To tackle this problem, we used a shifter network to correct the world camera video for eye movements (Walker et al., 2019; Yates et al., 2021). The shifter network takes as input the horizontal (theta) and vertical (phi) eye angle, along with the vertical head orientation (pitch) to approximate cyclotorsion (Wallace et al., 2013), and outputs the affine transformation for horizontal and vertical translation and rotation, respectively (Figure S2). We trained the shifter network and a GLM end-to-end with cross validation,to determine the camera correction parameters that best enable prediction of neural activity for each recording session (Figure 2A; see Methods). This analysis draws on the relatively large numbers of simultaneously recorded units as it determines the best shift parameters by maximizing fits across all neurons, thereby determining the general parameters of the eye camera to world camera transformation rather than being tailored to individual neurons.

**Figure 2:**
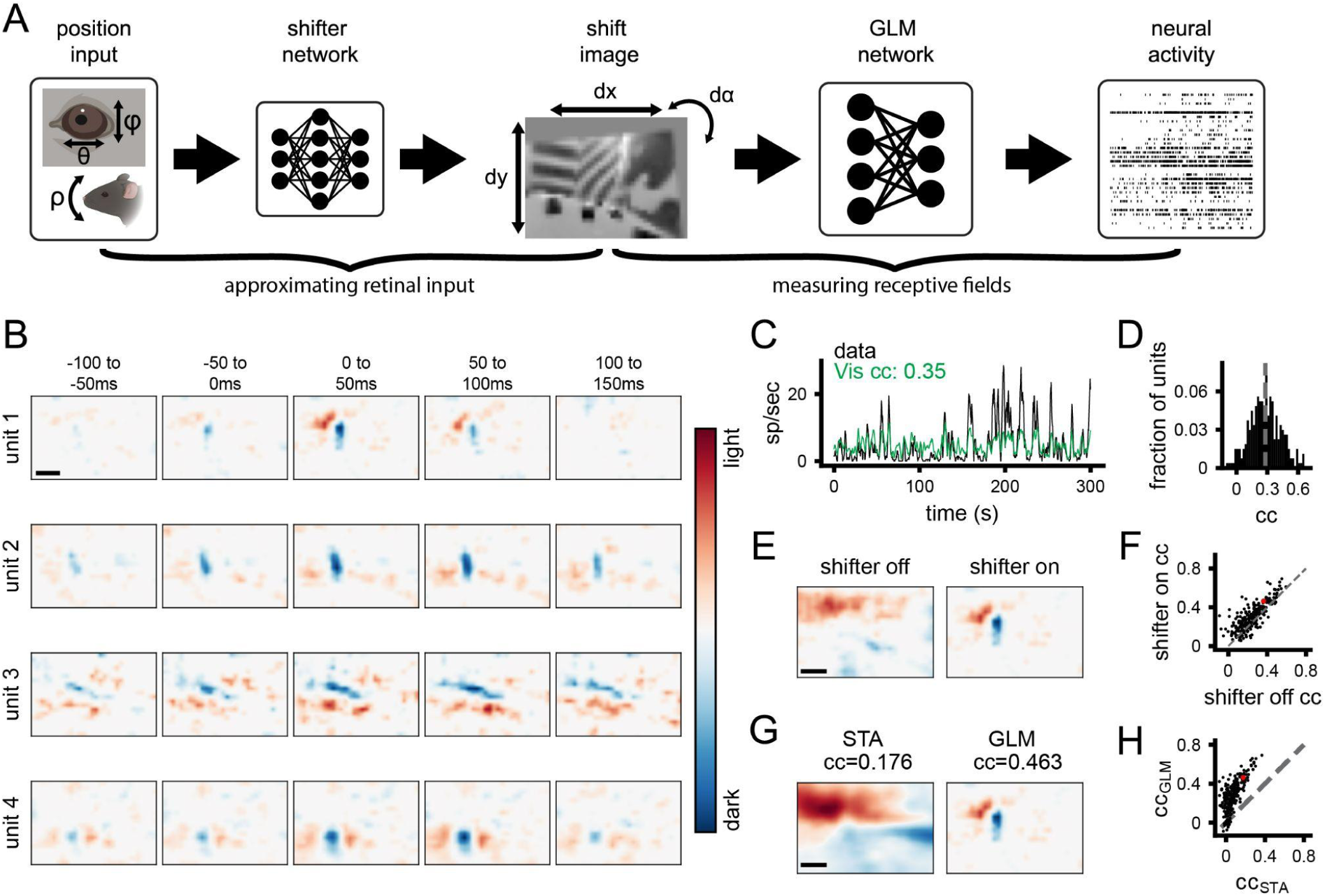
A generalized linear model accurately estimates spatiotemporal receptive fields during free movement. **A)** Schematic of processing pipeline. Visual and positional information is used as input into the shifter network, which outputs parameters for an affine transformation of the world-camera image. The transformed image frame is then used as the input to the GLM network to predict neural activity. **B)** Four example freely moving spatiotemporal visual receptive fields. Scale bar for RFs represents 10 degrees. **C)** Example actual and predicted smoothed (2 s window) firing rates for unit 3 in B. **D)** Histogram of correlation coefficients (cc) for the population of units recorded. Average cc shown as gray dashed line. **E)** Example of a freely moving RF with the shifter network off (left) and on (right) at time lag 0 ms. Colormap same as B. **F)** Scatter plot showing cc of predicted versus actual firing rate for all units with the shifter network off vs on. Red point is the unit shown in E. **G)** Example receptive field calculated via STA (left) versus GLM (right). **H)** Scatter plot showing cc of predicted vs actual firing rate for all units, as calculated from STA or GLM. Red point is the unit shown in G.

The outputs of the shifter network (Figure S2A-C) show that it converts the two axes of eye rotation (in degrees) into a continuous and approximately orthogonal combination of horizontal and vertical shifts of the worldcam video (in pixels), as expected to compensate for the alignment of the horizontal and vertical axes of the eye and world cameras. These outputs were also consistent in cross-validation across subsets of the data (coefficient of determination R^2^, dx=0.846, dy=0.792, da=0.945; Figure S2A-C). When the shifts were applied to the raw world camera video it had the qualitative effect of stabilizing the visual scene in between rapid gaze shifts, as would be expected from the vestibulo-ocular reflex and “saccade-and-fixate” eye movement pattern described previously in mice (Video S2; (Meyer et al., 2020; Michaiel et al., 2020). We quantified this by computing the total horizontal and vertical displacement of the raw and shifted world camera video based on image registration between sequential frames. When corrected for eye position, continuous motion of the image is converted into the step-like pattern of saccade-and-fixate (Figure S2D) and the image is stabilized to within 1 deg during the fixations (Figure S2E,F; (Michaiel et al., 2020). This eye-corrected retinocentric image was then used as input for the GLM network to predict neural activity in subsequent analysis.

We estimated spatiotemporal RFs during free movement using a GLM to predict single-unit activity from the corrected world camera data. Single-unit RFs measured during free movement had clear on and off sub-regions and a transient temporal response (Figure 2B). To our knowledge, these are the first visual receptive fields measured from a freely moving animal. It should be noted that the temporal response is still broader than would be expected, which likely reflects the fact that the GLM cannot fully account for strong temporal correlations in the visual input. Furthermore, the GLM predicted the continuous time-varying firing rate of units during free movement (Figure 2C). Across the population of neurons recorded (N=268 units, 4 animals), neural activity predicted from the corrected world camera data was correlated with the actual continuous firing rate (CC mean 0.28, max 0.69; Figure 2D). These values are on par with those obtained from mapping V1 RFs in awake and anesthetized head-fixed animals (Carandini et al., 2005).

To demonstrate the impact of correcting the visual input for eye movements, we computed RFs from the raw, uncorrected world camera data. This resulted in single-unit RFs becoming blurred, and reduced the ability to predict neural activity (Figure 2E,F; shifter on vs. off p=8.17e-23, paired t-test). Nonetheless, it is notable that the overall improvement was modest (mean increase in cc=0.06) and although some units required the shifter network, many units maintained a similar ability to predict firing rate even without the shifter. This is perhaps due to the large size of receptive fields relative to the amplitude of eye movements in the mouse (see Discussion). To determine the relative benefit of the GLM approach relative to a simpler reverse correlation spike-triggered average (Chichilnisky, 2001), we compared receptive fields and ability to predict firing rate from these two methods (Figure 2G-H).

Receptive fields from the STA were much broader and appeared to reflect structure from the environment (Figure 2G), as expected since the STA will not account for spatiotemporal correlations in the input. Correspondingly, the STA performed much worse than the GLM in predicting neural activity (Figure 2H; p=2e-93). Finally, as an additional verification that the GLM method is able to accurately reconstruct RFs from limited data and that natural scene statistics are not biasing the RF estimates, we simulated neural activity based on Gabor RFs applied to the world camera data. The results demonstrate that the GLM can reconstruct simulated RFs with high accuracy, resulting in reconstructed RFs that are both qualitatively and quantitatively similar to the original (Figure S2F,G).

### Comparison of receptive fields measured under freely moving versus head-fixed conditions

To determine whether RFs measured during free movement were comparable to those measured using traditional visual physiology methods, we compared them to RFs measured using a white noise stimulus under head-fixed conditions. The large majority of units were active (mean rate >1Hz) during each of these conditions (Figure 3A) and in each condition roughly half the units had a fit that significantly predicted neural activity, with slightly more in freely moving (Figure 3A). Overall, many neurons that had a clear white noise RF also had a clear RF from freely moving data (Figure 3B), which closely matched in spatial location, polarity, and number of sub-regions. To quantitatively compare RFs, we calculated the pixel-wise correlation coefficient between them. To provide a baseline for this metric, we first performed a cross validation test-retest by comparing the RFs from the first and second half of each recording separately (Figure S3). The mean test-retest cc was 0.46 for head-fixed and 0.58 and freely moving. We considered a unit to have a robust test-retest RF if this pixel-wise cc was greater than 0.5 (Figure S3C), and then evaluated the similarity of RFs for units that had robust fits in both conditions. The distribution of correlation coefficients between head-fixed and freely moving RFs for these units (Figure 3C) shows a strong correspondence for RFs across the two conditions (Figure 3C; 74% of units had a significant cc versus shuffled data). Taken together, these results show that for the units that had clearly defined RFs in both conditions, RFs measured with freely moving visual physiology are similar to those measured using traditional methods, despite the dramatically different visual input and behavior between these two conditions.

**Figure 3:**
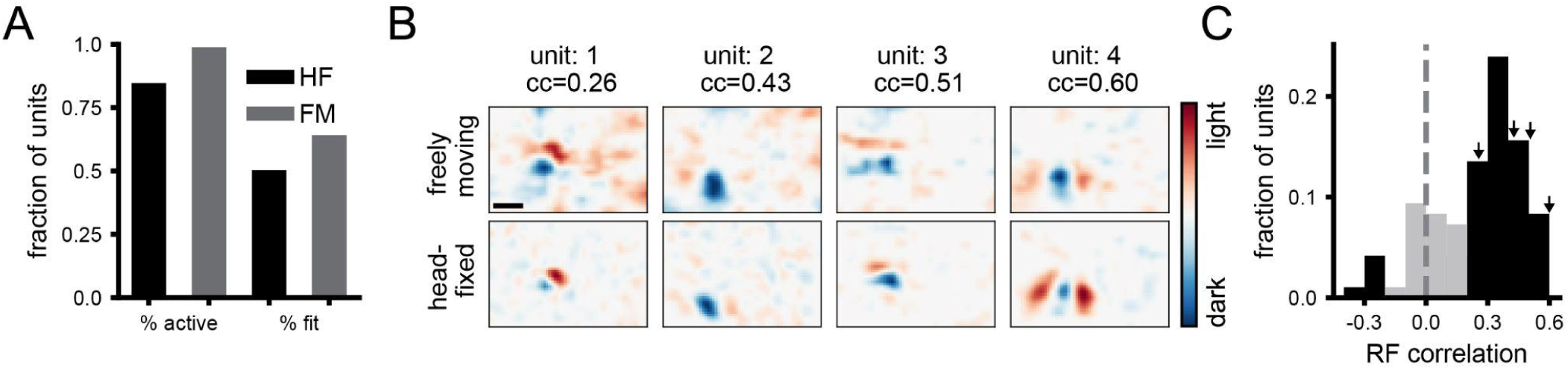
Comparison of receptive fields measured under freely moving versus head-fixed conditions. **A)** Fraction of units that were active (>1 Hz firing rate) and that had significant fits for predicting firing rate, in head-fixed and freely moving conditions. **B)** Example spatial receptive fields measured during free movement (top) and using a white noise mapping stimulus while head-fixed (bottom) at time lag 0 ms. Scale bar in top left is 10 deg. **C)** Histogram of correlation coefficients between freely moving and head-fixed RFs. Black color indicates units that fall outside two standard deviations of the shuffle distribution. Arrows indicate locations in the distribution for example units in A.

### V1 integrates visual and position signals

We next sought to determine whether and how eye/head position modulate V1 neural activity during free movement, based on measurement of pupil position from the eye camera and head orientation from the IMU. Strikingly, many single units showed tuning for eye position and/or head orientation, with 25% (66/268) of units having a modulation index [MI; (rate_max_-rate_min_)/(rate_max_+rate_min_)] greater than 0.33 for at least one position parameter, which equates to a two-fold change in firing rate (Figure 4A-C). To determine whether single-unit activity was better explained by visual input or eye/head position, we fit GLMs using either one as input. For most units (189/268 units, 71%), firing rate was better explained by a visual model, although the activity of some units was better explained by eye/head position (Figure 4D,E; 78/268 units, 29%). It should be noted that the units that were better fit by position model might nonetheless be better described by a more elaborate visual model.

**Figure 4:**
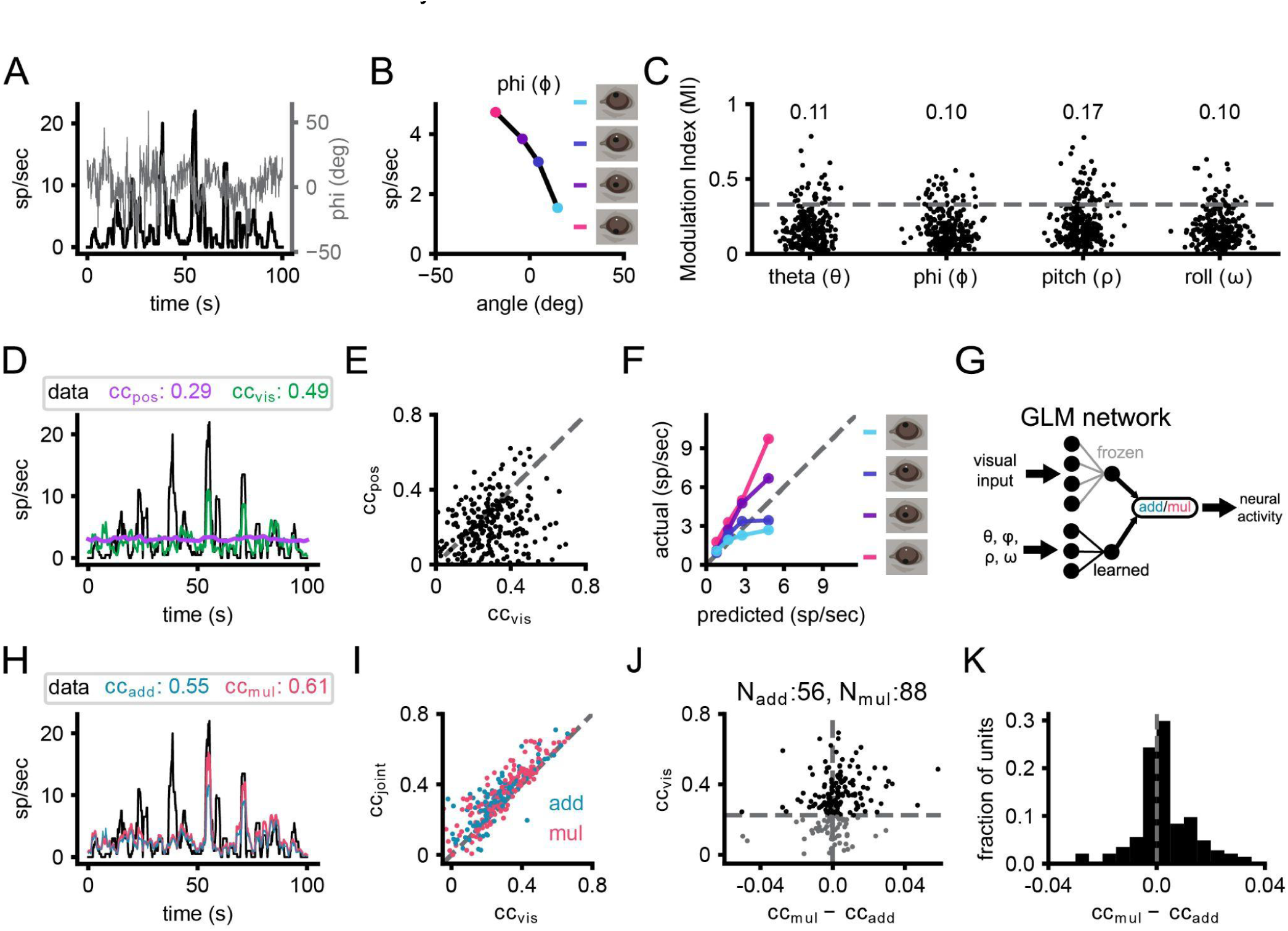
V1 neurons integrate visual and position signals. **A)** Overlay of vertical eye angle (phi; gray) and the smoothed firing rate of an example unit (black). **B)** Example tuning curve for head pitch. Colored points denote the quartiles of phi corresponding to panel F. **C)** Scatter of the modulation indices for eye position and head orientation (N=268 units, 4 animals). Numbers at top of the plot represent the fraction of units with significant tuning. **D)** Same unit as A. Example trace of smoothed firing rates from neural recordings and predictions from position-only and visual-only fits. **E)** Scatter plot of cc for position-only and visual-only fits for all units. **F)** Gain curve for the same unit in A and C. Modulation of the actual firing rates based on phi indicated by color. **G)** Schematic of joint visual and position input training. Visual spatiotemporal weights are frozen and position weights are learned. Output of visual and position modules are either added or multiplied together to predict firing rates. **H)** Same unit as A, C, and E. Smoothed traces of the firing rates from the data, additive and multiplicative fits. **I)** Correlation coefficient for visual-only versus joint fits. Each point is one unit, color coded for the joint fit that performed best. **J)** Comparison of additive and multiplicative fits for each unit. Units characterized as multiplicative are to the right of the vertical dashed line, while additive ones are to the left. Horizontal dashed line represents threshold set for the visual fit, since in the absence of a predictive visual fit, a multiplicative modulation will be similar to an additive modulation. **K)** Histogram of the difference in cc between additive and multiplicative models. The visual threshold from I was applied to the data.

To gain a qualitative understanding of how V1 neurons might combine visual and position information, we plotted predicted firing rates from visual-only GLM fits against the actual firing rates binned into quartiles based on eye/head position (example in Figure 4F). While the data should lie on the unity line in the absence of position modulation, additive integration would shift the entire curve up or down, and multiplicative integration would cause a slope change. Across the population of recorded neurons, many units showed evidence of gain modulation that tended to appear more multiplicative than additive.

To directly quantify the integration of visual and eye/head position information, and in particular to test whether this was additive or multiplicative, we trained two additional models: additive and multiplicative joint-encoding of visual and position information. To train the joint fit of visual and position signals, we froze the weights of the initial visual fits and trained positional weights that either added to or multiplied the visual signal for each unit (Figure 4G). Incorporating eye position and head orientation enables the model to more accurately predict large changes in the firing rate (Figure 4H). The inclusion of positional information almost universally improved predicted neural activity compared to visual fits alone (Figure 4I). For units that had a significant visual fit (cc>0.22, cross-validated, N=173 units), incorporating positional information resulted in an average fractional increase in correlation of 34% (0.07 average increase in cc). Multiplicatively combining visual and positional signals generated predictions that more closely matched actual firing rates than an additive combination in a majority of units (Figure 4J,K; p=0.0005, one sample t-test cc_mult_-cc_add_ for units with significant visual-only fits versus gaussian distribution with mean=0), suggesting visual and position signals in mouse V1 are more often integrated nonlinearly, consistent with previous studies in primate visual and parietal cortex (Andersen and Mountcastle, 1983; Morris and Krekelberg, 2019).

To further characterize the head and eye position modulations, we performed additional experiments recording V1 activity during free movement in nearly total darkness, followed by recording in the standard light condition. A significant fraction of neurons were modulated by at least 2:1 in the dark (Figure S4A,B; dark: 17%, 41/241; light: 31%, 75/241 units). Comparing the degree of modulation in the light vs dark for individual units revealed that the degree of tuning often shifted (Figure S4C), with some increasing their position tuning (consistent with an additive modulation that has a proportionally larger effect in the absence of visual drive) and others decreasing their position tuning (consistent with a multiplicative modulation that is diminished in the absence of a visual signal to multiply). In addition, to test whether position modulation might result from the abrupt transition from head-fixed recordings to free movement, we compared the degree of modulation during the first and second half of free movement sessions, and found no consistent change (Figure S4D). Finally, to test whether there was a bias in tuning for specific eye/head positions (e.g., upward versus downward pitch), we examined the weights of the position fits, which showed distributions centered around zero (Figure S4E), indicating that tuning for both directions was present for all position parameters, across the population.

Many response properties have been shown to vary across the cell types and layers of mouse V1 (Niell and Scanziani, 2021). Separating recorded units into putative excitatory and inhibitory, based on spike waveform as performed previously (Niell and Stryker, 2008), demonstrated that the visual fit performed better than than head/eye position for putative excitatory neurons, while the contributions were roughly equal for putative inhibitory cells (Figure S4F). This may be explained by the fact that putative excitatory neurons in mouse V1 have more linear visual responses (Niell and Stryker, 2008). We also examined whether the contribution of visual versus position information varied by laminar depth, and found no clear dependence (Figure S4G,H).

Finally, we examined the role of two factors that are known to modulate activity in mouse V1: locomotor speed and pupil diameter (Niell and Stryker, 2010; Reimer et al., 2014; Vinck et al., 2015). It is important to note that our GLM analysis excludes periods when the head is completely still, since that leads to dramatic over-representation of specific visual inputs that presents a confound in fitting the data. Therefore, the results presented above do not include the dramatic shift from non-alert/stationary to alert/moving that has been extensively studied (McGinley et al., 2015). Nonetheless, we find that including speed and pupil in the fit does indeed predict a part of the neural activity (Figure S4I). However this does not occlude the contribution from head/eye position or visual input. Examination of the weights in a joint fit of all parameters together demonstrates that although the contribution of locomotor speed is greater than any one individual position parameter (Figure S4J), the summed weights of head/eye position parameters are still the largest contribution (Figure S4K). It is also interesting to note that although head and eye position are often strongly correlated in the mouse due to compensatory eye movements (Meyer et al., 2020; Michaiel et al., 2020), the weights for each of these parameters (Figure S4J) are roughly equal in the GLM fit that can account for these correlations, demonstrating that both head and eye may contribute independently to coding in V1.

## DISCUSSION

Nearly all studies of neural coding in vision have been performed in subjects that are immobilized in some way, ranging from anesthesia to head and/or gaze fixation, which greatly limits the ability to study the visual processing that occurs as an animal moves through its environment. One important component of natural movement is the integration of the incoming visual information with one’s position relative to the scene. In order to determine how individual neurons in mouse V1 respond to visual input and eye/head position, we implemented an integrated experimental and model-based data analysis approach to perform visual physiology in freely moving mice. Using this approach, we demonstrate the ability to estimate spatiotemporal visual receptive fields during free movement, show that individual neurons have diverse tuning to head and eye position, and find that these signals are often combined through a multiplicative interaction.

### Integration of visual input and eye/head position

The ongoing activity of many units in V1 was modulated by both eye position and head orientation, as demonstrated by empirical tuning curves (Figure 4B) and model-based prediction of neural activity based on these parameters (Figure 4D). Modulation of neural activity in V1 and other visual areas by eye position (Andersen and Mountcastle, 1983; Durand et al., 2010; Rosenbluth and Allman, 2002; Trotter and Celebrini, 1999; Weyand and Malpeli, 1993) and head orientation (Brotchie et al., 1995; Guitchounts et al., 2020b) has been observed across rodents and primates, and fMRI evidence suggests human V1 encodes eye position (Merriam et al., 2013). Many of the position-tuned units we observed were also visually responsive, with clear spatiotemporal receptive fields.

In order to determine how these position signals were integrated with visual input, we used the GLM model trained on visual input only and incorporated either an additive or multiplicative signal based on a linear model of the eye/head position parameters. For neurons that had both a significant visual and position component, we found that the majority were best described by a multiplicative combination. This multiplicative modulation corresponds to a gain field, a fundamental basis of neural computation (Salinas and Abbott, 1996; Salinas and Sejnowski, 2001). Gain fields have been shown to serve a number of roles, including providing an effective mechanism for coordinate transformations as they enable direct readout of additive or subtractive combinations of input variables, such as the transformation from retinotopic to egocentric position of a visual stimulus. Studies in head-fixed primates have demonstrated gain fields for eye position (Andersen and Mountcastle, 1983; Morris and Krekelberg, 2019; Salinas and Sejnowski, 2001) and head orientation (Brotchie et al., 1995), and similar gain modulation for other factors such as attention (Salinas and Abbott, 1997). The demonstration of gain modulation by eye/head position in freely moving mice shows that this mechanism is engaged under natural conditions with complex movement.

Given the presence of gain fields in mouse visual cortex, two immediate questions arise: what are the sources of the position signals, and what are the cellular/circuit mechanisms that give rise to the gain modulation? Regarding sources, evidence suggests eye position signals arrive early in the visual system, perhaps even at the level of the thalamic lateral geniculate nucleus (Lal and Friedlander, 1990), while head orientation information could be conveyed through secondary motor cortex (Guitchounts et al., 2020a) retrosplenial cortex (Vélez-Fort et al., 2018) or from neck muscle afferents (Crowell et al., 1998). Regarding the mechanism, multiplicative interactions have been suggested to arise from synaptic interactions including active dendritic integration, recurrent network interactions, changes in input synchrony, balanced excitatory/inhibitory modulatory inputs, and classic neuromodulators (Salinas and Abbott, 1996; Salinas and Sejnowski, 2001; Silver, 2010). Future research could take advantage of genetic methods available in mice to determine the neural circuit mechanisms that implement this computation (Luo et al., 2018; Niell and Scanziani, 2021; O’Connor et al., 2009).

This multiplicative interaction can also be viewed as a form of nonlinear mixed selectivity, which has been shown to greatly expand the discriminative capacity of a neural code (Nogueira et al., 2021; Rigotti et al., 2013). The implications of nonlinear mixed selectivity have primarily been explored in the context of categorical variables, rather than continuous variables as observed here. In this context it is interesting to note that a significant number of units were nonetheless best described by an additive interaction. In an additive interaction the two signals are linearly combined, providing a factorized code where each signal can be read out independently. It may be that having a fraction of neurons using this linear interaction provides flexibility by which the visual input and position can be directly read out, along with the nonlinear interaction that allows computations such as coordinate transformations.

### Methodological considerations

We estimated the visual input to the retina based on two head-mounted cameras – one to determine the visual scene from the mouse’s perspective, and one to determine eye position and thereby correct the head-based visual scene to account for eye movements. Incorporation of eye position to correct the visual scene significantly improved the ability to estimate receptive fields and predict neural activity. Although head-fixed mice only make infrequent eye movements, freely moving mice (and other animals) make continual eye movements that both stabilize gaze by compensating for head movements and shift the gaze via saccades (Meyer et al., 2020; Michaiel et al., 2020). As a result, eye position can vary over a range of ±30 degrees (theta std: 16.5 deg, phi std: 17.8 deg in this study). Indeed, without eye movement correction many units did not have an estimated receptive field with predictive power (Figure 2F). Nonetheless, it is notable that some units were robustly fit even without correction – this likely reflects that fact that the eye is still within a central location a large fraction of the time (63% of timepoints within ±15 deg for theta, phi) and typical receptive fields in mouse V1 are on the order of 10-20 degrees (Niell and Stryker, 2008; Van den Bergh et al., 2010).

We estimated spatiotemporal receptive fields and predicted neural activity during free movement using a generalized linear model – a standard model-based approach in visual physiology (Pillow et al., 2008). Despite its simplicity – it estimates the linear kernel of a cell’s response – the GLM approach allowed us to estimate receptive fields in many neurons (39% of freely moving RFs significantly matched head-fixed white-noise RFs). These results are comparable to the fraction of units with defined STA receptive fields measured in head-fixed mice (64% of simple cells, 34% of total population in (Niell and Stryker, 2008); 49% of total population in (Bonin et al., 2011). The model fits were also able to predict a significant amount of ongoing neural activity (CC mean=0.29, max=0.73). Although this is still generally a small fraction of total activity, this is in line with other studies (Carandini et al., 2005; de Vries et al., 2020) and likely represents the role of additional visual features beyond a linear kernel, as well as other non-visual factors that modulate neural activity (Musall et al., 2019; Niell and Stryker, 2010; Stringer et al., 2019). A more elaborate model with nonlinear interactions would likely do a better job of explaining activity in a larger fraction of units; indeed, “complex” cells (Hubel and Wiesel, 1962) are not accurately described by a single linear kernel. However, for this initial characterization of receptive fields in freely moving animals, we chose to use the GLM since it is a well-established method, it is a convex optimization guaranteed to reach a unique solution, and the resulting model is easily interpretable as a linear receptive field filter. The fact that even such a simple model can capture many neurons’ responses both shows the robustness of the experimental approach, and opens up the possibility for the use of more elaborate and nonlinear models, such as multi-component (Butts, 2019) or deep neural networks (Bashivan et al., 2019; Ukita et al., 2019; Walker et al., 2019). Implementation of such models may require extensions to the experimental paradigm such as longer recording times to fit a greater number of parameters.

### Freely moving visual physiology

Visual neuroscience is dominated by the use of head-restrained paradigms, in which the subject cannot move through the environment. As a result, many aspects of how vision operates in the natural world remain unexplored (Leopold and Park, 2020; Parker et al., 2020). Indeed, the importance of movement led psychologist J. J. Gibson to consider the legs a component of the human visual system, and provided the basis for his ecological approach to visual perception (Gibson, 1979). The methods we developed here can be applied more broadly to enable a Gibsonian approach to visual physiology that extends beyond features that are present in standard head-fixed stimuli. While natural images and movies are increasingly used to probe responses of visual neurons in head-fixed conditions, these are still dramatically different from the visual input received during free movement through complex three-dimensional environments. This includes cues resulting from self-motion during active vision, such as motion parallax, loom, and optic flow that can provide information about the three-dimensional layout of the environment, distance, object speed, and other latent variables. Performing visual physiology in a freely moving subject may facilitate the study of the computations underlying these features.

Accordingly, a resurgent interest in natural behaviors (Datta et al., 2019; Dennis et al., 2021; Juavinett et al., 2018; Miller et al., 2022) provides a variety of contexts in which to study visual computations in the mouse. However, studies of ethological visual behaviors typically rely on measurements of neural activity made during head-fixation, rather than during the behavior itself (Boone et al., 2021; Hoy et al., 2019). Freely moving visual physiology is a powerful approach that ultimately can enable quantification of visual coding during ethological tasks to determine the neural basis of natural behavior.

## AUTHOR CONTRIBUTIONS

E.T.T.A., P.R.L.P., C.M.N. conceived the project. E.T.T.A. led design and implementation of computational analysis and P.R.L.P. led design and implementation of experiments. ESPL contributed to data collection. DMM contributed to data pre-processing. E.T.T.A. and P.R.L.P. generated figures. E.T.T.A., P.R.L.P., and C.M.N. wrote the manuscript.

## ACKNOWLEDGEMENTS

We thank Drs. Michael Beyeler, Michael Goard, Alex Huk, David Leopold, Jude Mitchell, Matt Smear and members of the Niell lab for comments on the manuscript. We thank Geordi Helmick and the University of Oregon Technical Sciences Administration for assistance with graphic and hardware design/production, Dr. Yuta Senzai for assistance with chronic electrode implants and drive design, and Jonny Saunders for assistance with sensor fusion analysis. This work was supported by NIH grants R34NS111669-01, R01NS121919-01, and UF1NS116377 (C.M.N.).

## KEY RESOURCE TABLE

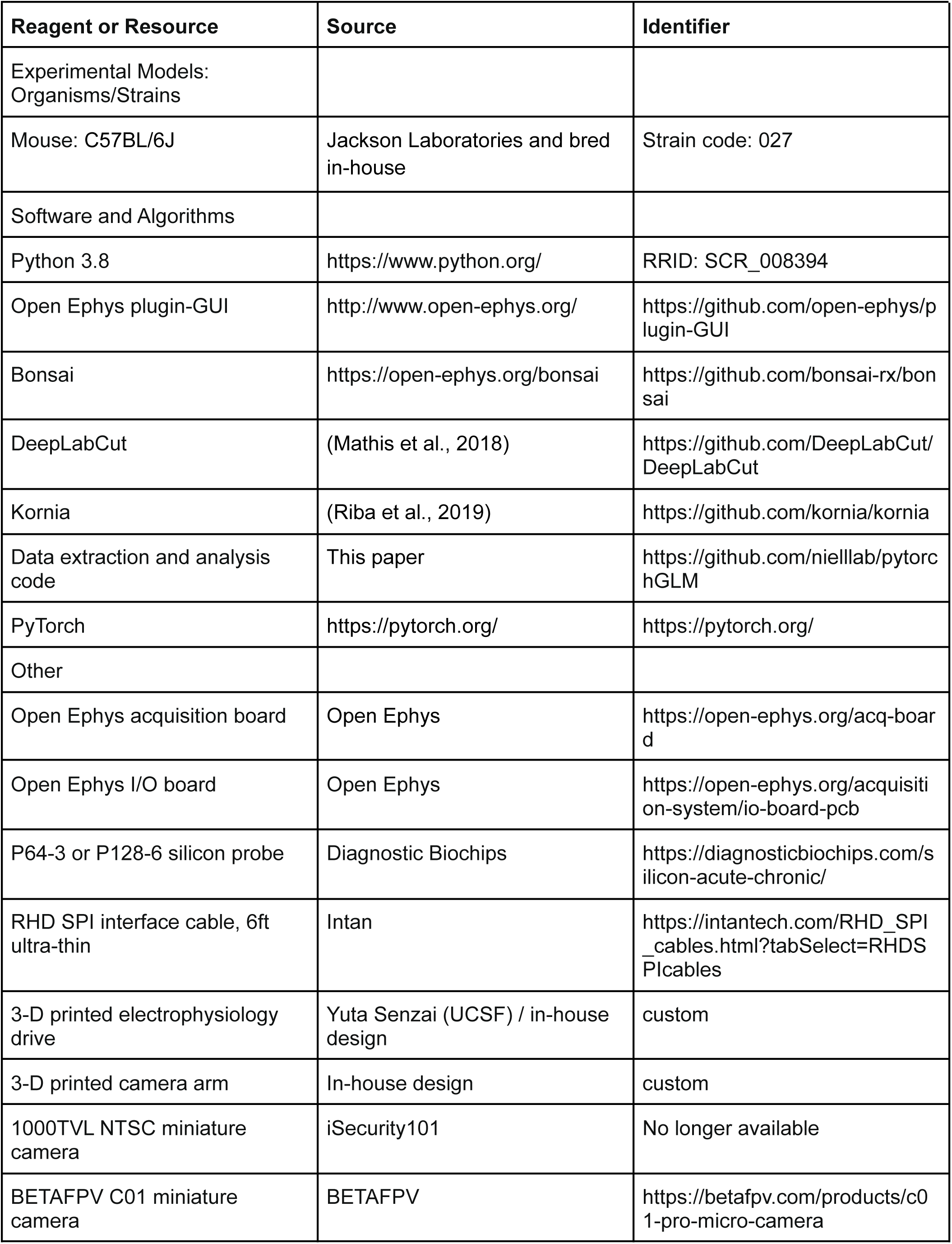
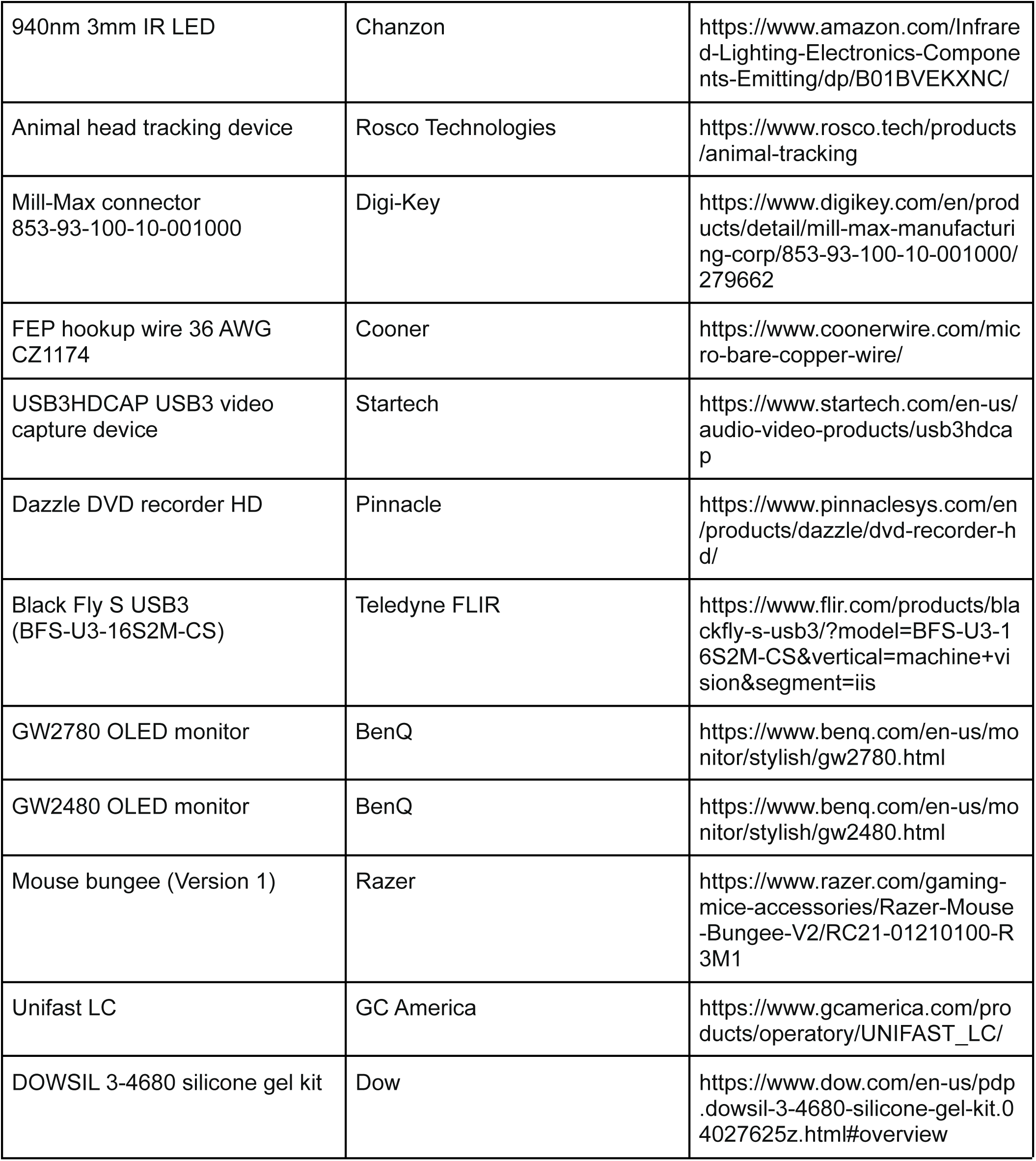

## METHODS

### Resource availability

#### Lead contact

Further information and requests for resources should be directed to and fulfilled by the lead contact, Dr. Cristopher M Niell.

#### Materials availability

This study did not generate new unique reagents.

#### Data and code availability

Data are available upon request. Code is available at: https://github.com/nielllab/pytorchGLM

### Experimental model and subject details

#### Animals

All procedures were conducted in accordance with the guidelines of the National Institutes of Health and were approved by the University of Oregon Institutional Animal Care and Use Committee. Three-to eight-month old adult mice (C57BL/6J, Jackson Laboratories and bred in-house) were kept on a 12 h light/dark cycle. In total, 4 female and 3 male mice were used for this study (head-fixed/freely moving: 2 females, 2 males; light/dark: 3 females, 2 males).

#### Surgery and habituation

Mice were initially implanted with a steel headplate over primary visual cortex to allow for head-fixation and attachment of head-mounted experimental hardware. After three days of recovery, widefield imaging (Wekselblatt et al 2016) was performed to help target the electrophysiology implant to the approximate center of left monocular V1. A miniature connector (Mill-Max 853-93-100-10-001000) was secured to the headplate to allow attachment of a camera arm (eye/world cameras and IMU; Michaiel et al., 2020). In order to simulate the weight of the real electrophysiology drive and camera system for habituation (6 g total), a ‘dummy’ system was glued to the headplate. Animals were handled by the experimenter for several days before surgical procedures, and subsequently habituated (~45 min) to the spherical treadmill and freely moving arena with hardware tethering attached for several days before experiments.

The electrophysiology implant was performed once animals moved comfortably in the arena. A craniotomy was performed over V1, and a linear silicon probe (64 or 128 channels, Diagnostic Biochips P64-3 or P128-6) mounted in a custom 3D-printed drive (Yuta Senzai, UCSF) was lowered into the brain using a stereotax to an approximate tip depth of 750 µm from the pial surface. The surface of the craniotomy was coated in artificial dura (Dow DOWSIL 3-4680) and the drive was secured to the headplate using light-curable dental acrylic (Unifast LC). A second craniotomy was performed above left frontal cortex, and a reference wire was inserted into the brain. The opening was coated with a small amount of sterile ophthalmic ointment before the wire was glued in place with cyanoacrylate. Animals recovered overnight and experiments began the following day.

#### Hardware and recording

The camera arm was oriented approximately 90 deg to the right of the nose and included an eye-facing camera (iSecurity101 1000TVL NTSC, 30 fps interlaced), an infrared-LED to illuminate the eye (Chanzon, 3 mm diameter, 940 nm wavelength), a wide-angle camera oriented toward the mouse’s point of view (BETAFPV C01, 30 fps interlaced) and an inertial measurement unit acquiring three-axis gyroscope and accelerometer signals (Rosco Technologies; acquired 30 kHz, downsampled to 300 Hz and interpolated to camera data). Fine gauge wire (Cooner, 36 AWG, #CZ1174CLR) connected the IMU to its control box, and each of the cameras to a USB video capture device (Pinnacle Dazzle or StarTech USB3HDCAP). A top-down camera (FLIR Blackfly USB3, 60 fps) recorded the mouse in the arena. The electrophysiology headstage (built into the silicon probe package) was connected to an OpenEphys acquisition system via an ultra thin cable (Intan #C3216). The electrophysiology cable was looped over a computer mouse bungee (Razer) to reduce the combined impact of the cable and implant. We first used the OpenEphys GUI (https://open-ephys.org/gui) to assess the quality of the electrophysiology data, then recordings were performed in Bonsai (Lopes et al., 2015) using custom workflows. System timestamps were collected for all hardware devices and later used to align data streams through interpolation.

During experiments, animals were first head-fixed on a spherical treadmill to permit measurement of visual receptive fields using traditional methods, then were transferred to an arena where they could freely explore. Recording duration was approximately 45 minutes head-fixed, and 1hr freely moving. For head-fixed experiments, a 27.5 in monitor (BenQ GW2780) was placed approximately 27.5 cm from the mouse’s right eye. A contrast-modulated white noise stimulus (Niell and Stryker, 2008) was presented for 15 min, followed by additional visual stimuli, and the mouse was then moved to the arena. The arena was approximately 48 cm long by 37 cm wide by 30 cm high. A 24 in monitor (BenQ GW2480) covered one wall of the arena, while the other three walls were clear acrylic covering custom wallpaper including black and white high- and low-spatial frequency gratings and white noise. A moving black and white spots stimulus (Piscopo et al., 2013) played continuously on the monitor while the mouse was in the arena. The floor was a gray silicone mat (Gartful) and was densely covered with black and white Legos. Small pieces of tortilla chips (Juanita’s) were lightly scattered around the arena to encourage foraging during the recording, however animals were not water or food restricted.

### Method details

#### Data preprocessing

Electrophysiology data were acquired at 30 kHz and bandpass filtered between 0.01 Hz and 7.5 kHz. Common-mode noise was removed by subtracting the median across all channels at each timepoint. Spike sorting was performed using Kilosort 2.5 (Steinmetz et al., 2021), and isolated single units were then selected using Phy2 (https://github.com/cortex-lab/phy) based on a number of parameters including contamination (<10%), firing rate (mean >0.5 Hz across entire recording), waveform shape, and autocorrelogram. Electrophysiology data for an entire session were concatenated (head fixed stimulus presentation, freely moving period, or freely moving light and dark) and any sessions with apparent drift across the recording periods (based on Kilosort drift plots) were discarded. To check for drift between head-fixed and freely moving recordings, we compared the mean waveforms and noise level for each unit across the two conditions, based on a 2 ms window around the identified spike times in bandpass-filtered data (800-8000Hz). An example mean waveform, with its standard deviation across individual spike times, is shown in Figure S1A. To determine whether the waveform changed, indicative of drift, we calculated coefficient of determination (R^2^) between the two mean waveforms for each unit, which confirms a high degree of stability as the waveforms are nearly identical across conditions (Figure S1B). To determine whether the noise level changed, we computed the standard deviation across spike occurrences within each condition, for each unit (Figure S1C). There was no change in this metric between head-fixed and freely moving, indicating that there was not a change in noise level that might disrupt spike sorting in one condition specifically.

World and eye camera data were first deinterlaced to achieve 60 fps video. The world camera frames were then undistorted using a checkerboard calibration procedure (Python OpenCV), and downsampled to 30 by 40 pixels to reduce dimensionality and approximate mouse visual acuity. In order to extract pupil position from the eye video, eight points around the pupil were tracked with DeepLabCut (Mathis et al., 2018). We then fit these eight points to an ellipse and computed pupil position in terms of angular rotation (Michaiel et al., 2020). Sensor fusion analysis was performed on the IMU data (Jonny Saunders, University of Oregon) to calculate pitch and roll of the head. Pitch and roll were then passed through a median filter with window size 550 ms. All data streams were aligned to 50 ms bins through interpolation using system timestamps acquired in Bonsai.

#### GLM training

For all model fits, the data were partitioned into 10% groups, and were randomly sampled into cross-validation train and test sets (70%/30% split, respectively). Video frames were cropped by 5 pixels on each side to remove edge artifacts. Initially, a shifter network was trained on each recording session (see below) to estimate the appropriate horizontal shift, vertical shift, and rotation of the world camera video to correct for eye movements. The corrected eye camera data were then saved out and used for training. Eye and head position were z-scored and zero-centered before training and analysis. Four different networks were trained: 1) Eye position and head orientation signals only, 2) Visual input only, 3) Additive interaction between position and visual input, and 4) Multiplicative interaction between position and visual input. Units with a mean firing rate below 1 Hz in either head-fixed or freely moving were removed from the data set (17% of total units).

#### Network parameters

To train the model end-to-end and to speed up the computation we utilized the graphical processing unit (GPU) and pyTorch because the GLM is equivalent to a single-layer linear network. We then used a rectified linear activation function to approximate non-zero firing rates. Utilizing the GPU decreased training time for the model by multiple orders of magnitude (from over 500 hours down to 40 minutes for the entire dataset). L1 and L2 regularization was applied to the spatiotemporal filters of the visual model. The Adam optimization algorithm (Kingma and Ba, 2014) was used to update the parameters of the model to minimize prediction error. The loss and gradient of each neuron were computed independently in parallel so the full model represents the entire dataset. To account for the convergence of different parameters at different speeds as well as to isolate parameters for regularization, parameter groups were established within the optimizer with independent hyperparameters.

#### Shifter network

In order to correct the world camera video for eye movements, we trained a shifter network to convert eye position and torsion into an affine transformation of the image at each time point. For each recording session, eye angle and head pitch (theta, phi, and rho) were used as input into a feedforward network with a hidden layer of size 50, and output representing horizontal shift, vertical shift, and image rotation. The output of the network was then used to perform a differentiable affine transformation (Riba et al., 2019) to correct for eye movements. Head pitch was used as a proxy of eye torsion (Wallace et al., 2013), and eye position was zero-centered based on the mean position during the freely moving condition. The transformed image was then used as input into the GLM network to predict the neural activity. The shifter network and GLM were then trained together to minimize the error in predicted neural activity. During the shifter training (2000 epochs) no L1 regularization was applied to ensure a converged fit. Horizontal and vertical shift was capped at 20 pixels and rotation was capped at 45 deg. The eye corrected videos were saved out to be used for the model comparison training. The shifter network was trained on freely moving data, since eye movements are greatly reduced during head-fixation, but was applied to both head-fixed and freely moving data to align receptive fields across the two conditions.

#### Tuning and gain curves

Tuning curves for eye and head position were generated by binning the firing rates into quartiles so the density of each point is equal and then taking the average. For each gain curve we collected the time points of the firing rates that were within each quartile range for eye and head position, averaged the firing rates and then compared them with the predicted firing rates from the visual-only model. Each curve therefore represents how much each unit’s actual firing rate changed on average when the mouse’s eye or head was in the corresponding position.

#### Position-only model fits

Eye and head position signals were used as input into a single-layer network where the input dimension was four and the output dimension was the number of neurons. No regularization was applied during training due to the small number of parameters needed for the fitting. The learning rate for the weights and biases was 1e-3.

#### Visual-only model fits

Eye corrected world camera videos were used as input into the GLM network. The weights from the shifter training for each neuron were used as the initialization condition for the weights, while the mean firing rates of the neurons were used as the initialization for the biases. Parameters for the model were fit over 10,000 epochs with a learning rate of 1e-3. To prevent overfitting, a regularization sweep of 20 values log-base 10 distributed between 0.001 to 100 was performed. The model weights with the lowest test error were selected for each neuron.

#### Joint visual-position model fits

After the visual-only fits, the spatiotemporal weights and biases were frozen. A position module was then added to the model for which the input was the eye and head position signals (see Figure 4G). The output of the visual module was then combined with output of the position module in either an additive or multiplicative manner, then sent through a ReLu nonlinearity to approximate firing rates. The parameters for the position module were then updated with the Adam optimizer with learning rate 1e-3.

#### Speed and pupil diameter fits

To test the contribution of the speed and pupil diameter, the data were first z-scored and GLM fits were conducted with only speed and pupil, with eye/head position only and with speed, pupil and eye/head position. The explained variance (r^2^) of the predicted and actual firing rate was calculated between these models to show how these parameters contribute uniquely and sublinearly to the GLM fits. Additionally, we trained the joint fits with eye/head position and speed and pupil and calculated the total contribution of eye/head position versus speed and pupil (Figure S4I-K).

#### Post-training analysis

To better assess the quality of fits, the actual and output firing rates were smoothed with a boxcar filter with a 2 s window. The correlation coefficient (cc) was then calculated between smoothed actual and predicted firing rates of the test dataset. The modulation index of neural activity by position was calculated as the (max-min)/(max+min) of each signal. In order to distinguish between additive and multiplicative models (Figure 4J,K), a unit needs to have a good positional and visual fit. As a result, units which had an cc value below 0.22, or did not improve with incorporating position information were thresholded out for the final comparison.

#### Simulated RF reconstruction

We tested the ability of our GLM approach to recover accurate receptive fields using simulated data. Simulated RFs were created based on Gabor functions and applied to the eye movement-corrected world camera video as a linear filter to generate simulated neural activity, scaled to empirically match the firing rates of real neurons with an average firing rate of 14 Hz. The output was then passed through a Poisson process to generate binned spike counts. Using these simulated data, we then followed the same analysis as for real data to fit a visual GLM model and estimate RFs, using spatiotemporal weights set to zero for the initial conditions.

#### Test-retest analysis receptive fields

To assess how reliable the receptive fields were, we trained the GLM separately on the first and second half of each recording session. We then took the receptive fields that were mapped for each half and calculated the pixel-wise correlation coefficient (Figure S3). A threshold of 0.5 cc was then used as a metric for stable RFs within the same condition. The units that had a stable RF in both head-fixed and freely moving conditions were then used for the analysis in Figure 3.

#### Shifter controls and change in visual scene

Similar to the test-retest for receptive fields, we trained the shifter network on the first and second half of the data. Shifter matrices were created using a grid of eye and head angles after training to see how the network responds to different angles. The coefficient of determination (R^2^) was then calculated between the shifter matrices of the first and second half (Figure S2A-C). To further quantify the effect of the shifter network we used frame to frame image registration to measure the visual stability of the world camera video. Displacement between consecutive images was based on image registration performed with findTransformECC function in OpenCV. We computed the cumulative sum of shifts to get total displacement, then calculated standard deviation in the fixation intervals following analysis in (Michaiel et al., 2020).

#### Dark experiments and analysis

To eliminate all possible light within the arena, the entire behavioral enclosure was sealed in light-blocking material (Thorlabs BK5), all potential light sources within the enclosure were removed, and the room lights were turned off. Animals were first recorded in the dark (~20 min), then the arena lights and wall stimulus monitor were turned on (~20 min). As a result of the dark conditions, the pupil dilated beyond the margins of the eyelids, which made eye tracking infeasible. To counteract this, prior to the experiment, one drop of 2% Pilocarpine HCl Ophthalmic Solution was applied to the animal’s right eye to restrict the pupil to a size similar to that seen in the light. Once the pupil was restricted enough for tracking in the dark (~3 min) the animal was moved into the dark arena for recording, until the effects of the Pilocarpine wore off (~20 min), at which time the light recording began. Tuning curves for eye and head position were generated using the same method as in the light by binning the firing rates into quartiles so the density of each point is equal and then taking the average.

#### Quantification and statistical analysis

For shuffle distributions, we randomly shuffled spike times within the cross-validated train and test sets and then performed the same GLM training procedure. We defined significant values as two standard deviations away from the mean of the shuffle distribution. For paired t-tests, we first averaged across units within a session, then performed the test across sessions.

#### Figures

Some figure panels were generated using Biorender.com.

## Supplemental Figures

## Supplemental Videos

**Video S1:** Sample experimental data from a fifteen second period during free movement. Left: world camera video (top) and eye camera video (bottom). Right (from top): horizontal and vertical eye position (degrees), head orientation pitch and roll (degrees), and a raster plot of over 100 units (probe schematic bottom left). Vertical blue line in plots represents the current time. Note that the animal begins moving at ~4 s, accompanied by a shift in the dynamics of neural activity.

**Video S2:** Example of raw (left) and shifter network-corrected (right) world camera video from a single experiment. Note that the overall effect of the shifter network is to rotate and stabilize the image over short periods, interspersed by larger shifts in the image due to saccades.

---

**Figure S1:**
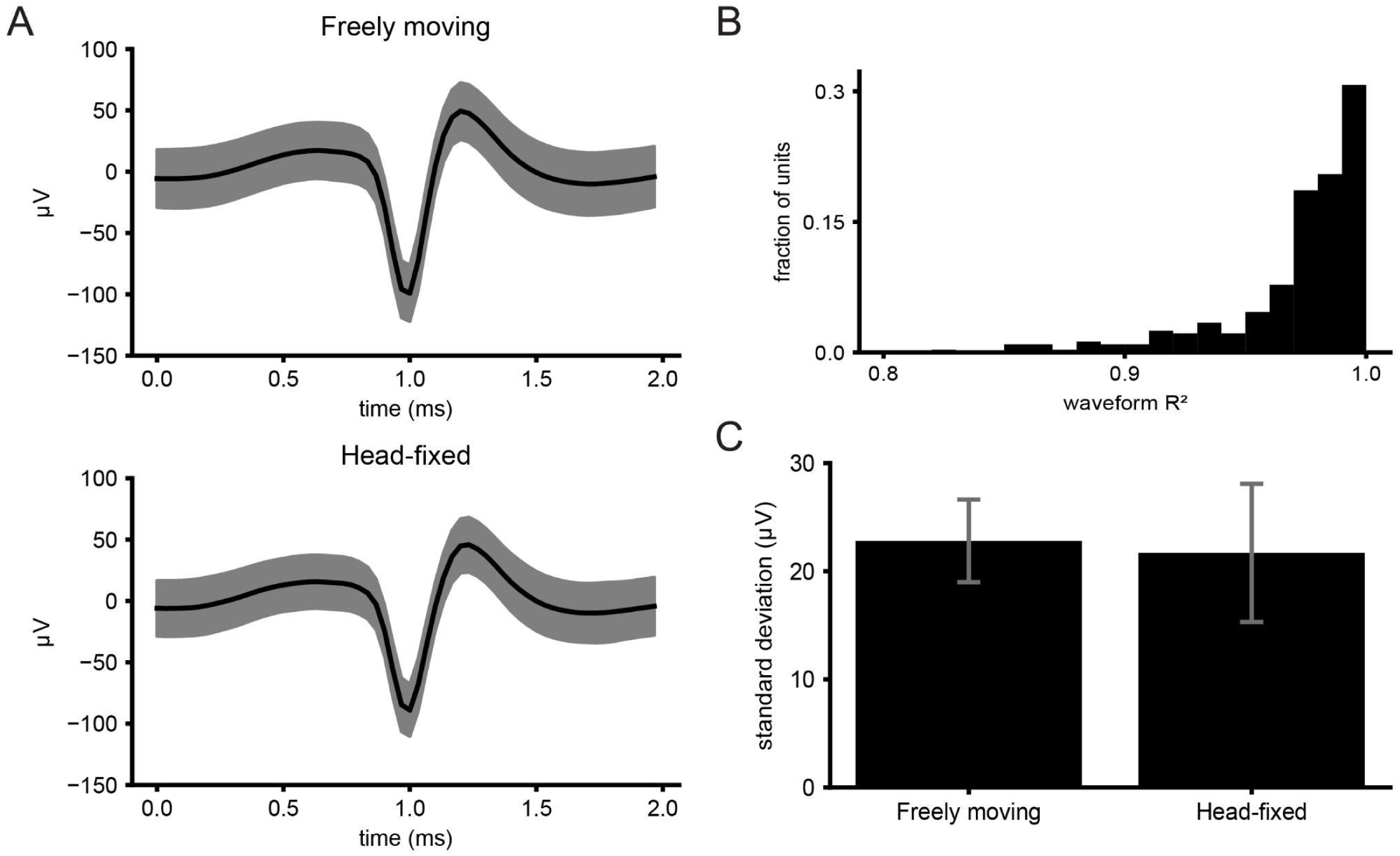
Comparison of spike waveforms across the experimental session. **A)** Top: Average spike waveform for one example unit in freely moving recording. Shaded region is one standard deviation. Bottom: Same unit as top but for head-fixed recording of the same unit in the same session. **B)** Histogram of coefficient of determination (R^2^) between units of freely moving and head-fixed recordings. **C)** Average standard deviation across 2 ms around spikes for freely moving (FM) and head-fixed (HF) recordings.

**Figure S2:**
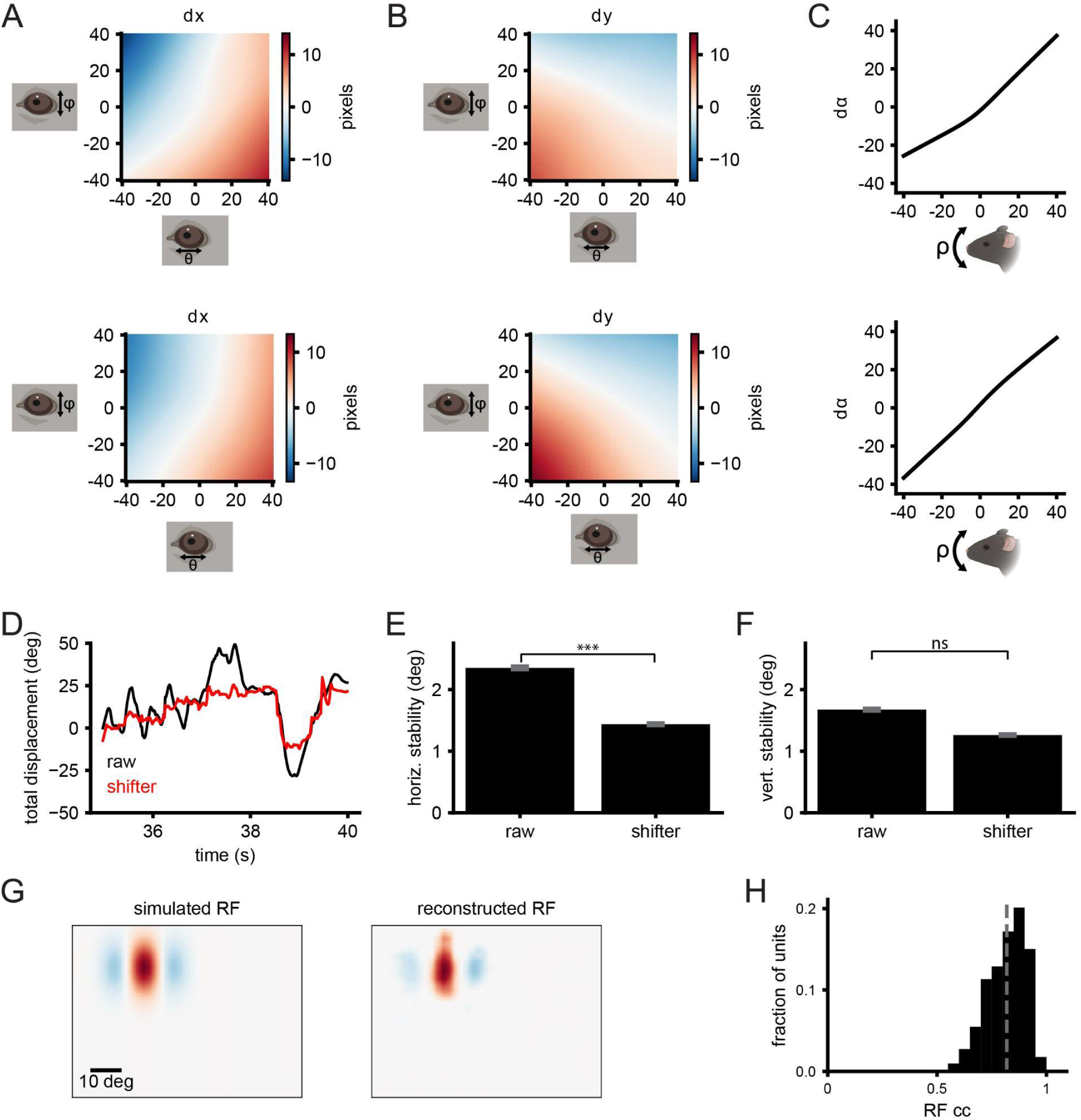
Quantification of shifter network performance. **A)** 2-d heat map of horizontal shift for values of theta and phi for first half (top) and second half (bottom) of example recording. **B)** Same as A but for the vertical shift of the image. **C)** Rotation of the image as a function of head pitch for the first half (top) and second half (bottom) of the recording. **D)** Image registration horizontal displacements for shifted and raw world camera video. **E)** Bar plot showing the average horizontal stability of visual angle for compensatory eye movements. **F)** Same as E but for vertical shifts. (***: p-value<0.0013) **G)** Simulated (left) and reconstructed (right) receptive fields with three sub-regions. Same training procedure as Figure 2A. **H)** Histogram of correlation coefficients between simulated and reconstructed RFs. The gray dashed line represents the mean of the distribution.

**Figure S3:**
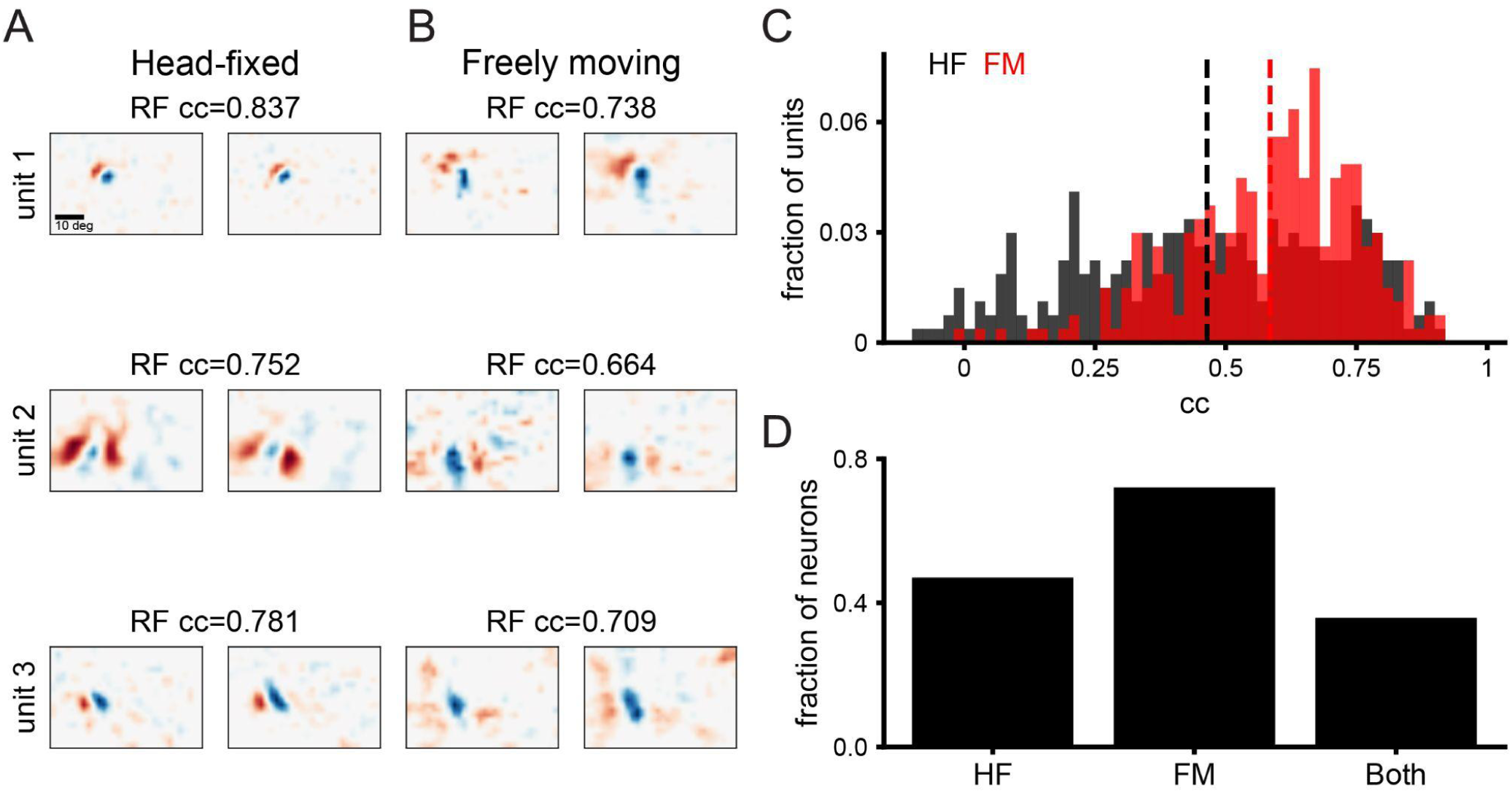
Test-retest analysis of receptive fields within head-fixed and freely moving recordings. **A)** Three example receptive fields mapped in the first (left) and second (right) half of a head fixed recording. Correlation coefficient (cc) given is the pixel-wise cc of the receptive fields. **B)** Same as A but for freely moving recording. **C)** Histogram of cc of receptive fields for first versus second half of recording for head-fixed (gray) and freely moving (red) conditions. Dashed lines indicate the mean of the distribution. **D)** Bar plot showing the fraction of units that have a significant cc between the first and second half of the recordings (cc>0.5).

**Figure S4:**
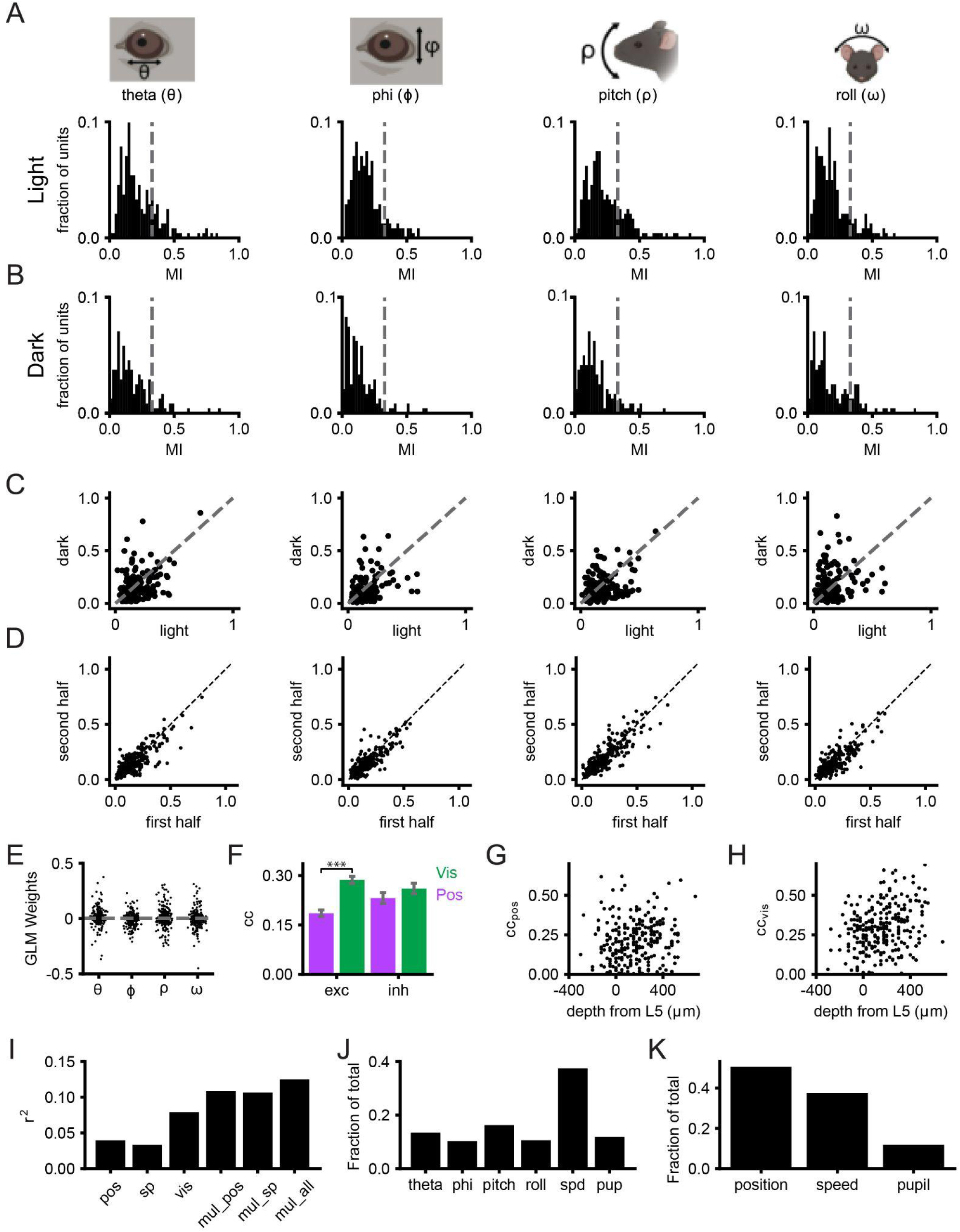
Eye/head position modulation in darkness, as a function of cell type/layer, and compared to pupil and locomotor speed. **A-D)** Columns correspond to analyses for theta, phi, pitch and roll respectively. **A)** Histograms of modulation index for single units recorded during free movement in the light. **B)** Same as A but recorded during free movement in darkness. C) Scatter plot comparison of light and dark modulation index for each unit. **D)** Modulation index calculated for first half and second half of the freely moving experiments in the light. **E)** Distribution of weights for position only GLM fit for eye/head position. **F)** Correlation coefficient (cc) of predicted versus actual firing rate for visual and position fits split by putative excitatory and inhibitory units. Error bars indicate standard error (***: p-value<0.001, between excitatory visual and position fits). **G)** Correlation coefficient as a function of depth from layer 5 for position fits (>0 deeper, <0 shallower). **H)** Same as G but for visual fits. **I)** Explained variance (r^2^) for position only (pos), speed and pupil only (sp), visual only (vis), multiplicative with eye/head position (mul_pos), multiplicative with speed and pupil (mul_sp), and multiplicative with eye/head position, speed and pupil (mul_all). **J)** The fraction of contribution of the weights for multiplicative fits with eye/head position, speed (spd) and pupil (pup). **K)** Same as J but summing together the contribution for eye/head position.

